# BioKEEN: A library for learning and evaluating biological knowledge graph embeddings

**DOI:** 10.1101/475202

**Authors:** Mehdi Ali, Charles Tapley Hoyt, Daniel Domingo-Fernández, Jens Lehmann, Hajira Jabeen

**Affiliations:** Rheinische Friedrich-Wilhelms-Universität Bonn, Bonn 53113, Germany; Department of Bioinformatics, Fraunhofer Institute for Algorithms and Scientific Computing (SCAI), Sankt Augustin 53754, Germany; Department of Enterprise Information Systems, Fraunhofer Institute for Intelligent Analysis and Information Systems (IAIS), Sankt Augustin 53754, Germany

## Abstract

Knowledge graph embeddings (KGEs) have received significant attention in other domains due to their ability to predict links and create dense representations for graphs’ nodes and edges. However, the software ecosystem for their application to bioinformatics remains limited and inaccessible for users without expertise in programming and machine learning. Therefore, we developed BioKEEN (Biological KnowlEdge EmbeddiNgs) and PyKEEN (Python KnowlEdge EmbeddiNgs) to facilitate their easy use through an interactive command line interface. Finally, we present a case study in which we used a novel biological pathway mapping resource to predict links that represent pathway crosstalks and hierarchies.

**Availability:** BioKEEN and PyKEEN are open source Python packages publicly available under the MIT License at https://github.com/SmartDataAnalytics/BioKEEN and https://github.com/SmartDataAnalytics/PyKEEN as well as through PyPI.

## 1. Introduction

Knowledge graphs (KGs) are multi-relational, directed graphs in which nodes represent entities and edges represent their relations (Bordes *et al.* 2013). While they have been successfully applied for question answering, information extraction, and named entity disambiguation outside of the biomedical domain, their usage in biomedical applications remains limited (Malone *et al.* 2018; Nickel *et al.,* 2016).

Because KGs are inherently incomplete and noisy, several methods have been developed for deriving or predicting missing edges (Nickel *et al.,* 2016). One is to apply reasoning based on formal logic to derive missing edges, but it usually requires a large set of user-defined formulas to achieve generalization. Another is to train knowledge graph embedding (KGE) models, which encode the nodes and relations in a KG into a low-dimensional, continuous vector-space that best preserves the structural characteristics of the KG (Wang *et al.,* 2017). These embeddings can be used to predict new relations between entities. In a biological setting, relation prediction not only enables researchers to expand their KGs, but also to generate new hypotheses that can be tested experimentally.

Here, we present BioKEEN (Biological KnowlEdge EmbeddiNgs): a Python package for training and evaluating KGEs on biological KGs that is accessible and facile for bioinformaticians without expert knowledge in machine learning through an interactive command line interface (CLI). Through the integration of the Bio2BEL software (https://github.com/bio2bel) within BioKEEN, numerous biomedical databases containing structured knowledge are directly accessible. Additionally, we have developed BioKEEN’s core component for training and evaluating KGE models, PyKEEN (Python KnowlEdge EmbeddiNgs), such that it can be reused in other domains.

While there exists other toolkits like OpenKE (Han et al., 2018) and scikit-kge (https://github.com/mnick/scikit-kge), they are not specialised for bioinformatics applications and they require more expertise in programming and in KGEs. To the best of our knowledge, BioKEEN is the first framework specifically designed to facilitate the use of KGE models for users in the bioinformatics community.

## 2. Software Architecture

The BioKEEN software package consists of three layers: 1) the model configuration layer, 2) the data acquisition and transformation layer, and 3) the learning layer (Figure 1).

**Figure 1.**
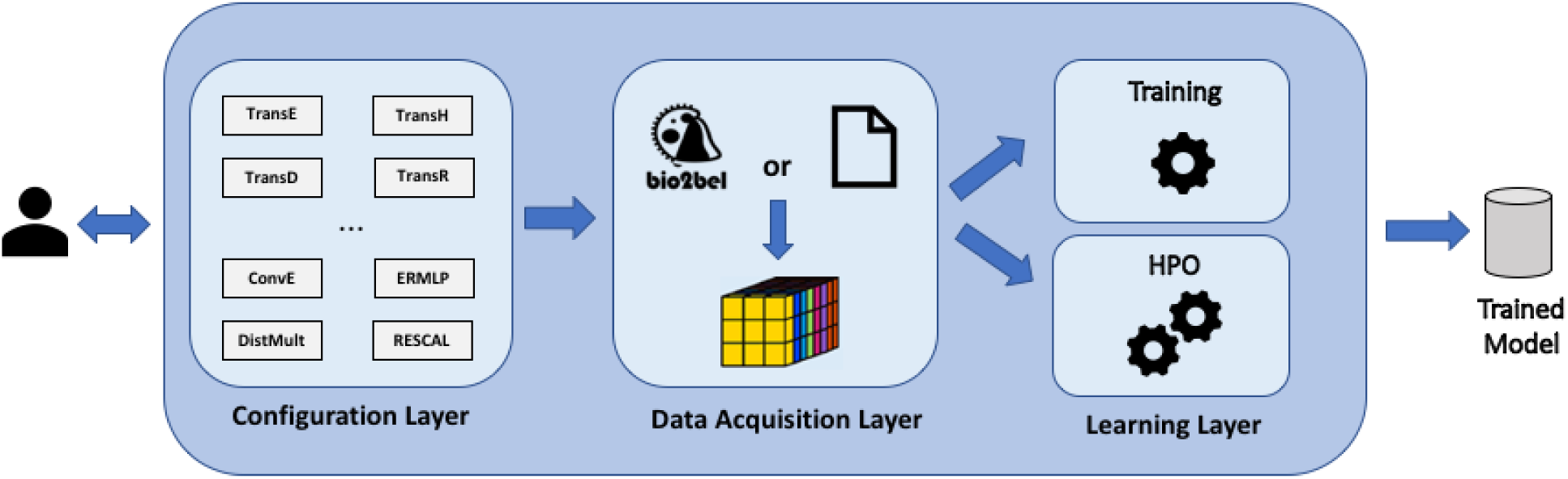
Software architecture of BioKEEN 1) **Configuration layer:** Users define experiments through the command line interface. 2) **Data acquisition layer:** Dataset(s) are (down-)loaded and transformed into a tensor. **3) Learning layer:** Model is trained with user-defined hyper-parameters or a hyper-parameter search is applied to find the best set of hyper-parameter values.

### 2.1 Configuration Layer

Because every KGE model has its own set of hyper-parameters, the configuration of an experiment for a non-expert can be very complicated and discouraging. This possible obstacle is addressed in the configuration layer through an interactive CLI that assists users in setting up their experiments (i.e., defining the datasets, the model, and its parameters)

Currently, we provide implementations of 10 embedding models (e.g., TransE, TransH, ConvE, etc. (Wang *et al., 2017*; Dettmers *et al.,* 2017)). A full list can be found in **Supplementary Table S1**. Moreover, BioKEEN can be executed in training and hyper-parameter optimization (HPO) mode.

### 2.2 Data Acquisition Layer

Because extracting and preparing training data can be a time-consuming process, BioKEEN integrates the Bio2BEL software to download and parse numerous biomedical databases (**Supplementary Table S2**). This allows users to focus on the experiments, to automatically incorporate the latest database versions, and to have access to new datasets as they are incorporated into Bio2BEL. In addition, users can provide their own datasets as tab-separated values, RDF, or from NDEx (Pratt *et al.*, 2015). BioKEEN processes the selected and provided datasets then transforms them into a tensor for further processing.

### 2.3 Learning Layer

Determining the appropriate values for the hyper-parameters of a KGE model requires both machine learning and domain specific knowledge. If the user specifies hyper-parameters, BioKEEN can be run directly in *training mode*. Otherwise, it first runs in *hyper-parameter optimization (HPO*) mode, where *random search* is applied to find suitable hyper-parameters values from (user) predefined sets. We implemented *random search* instead of the widely applied *grid search* because it converges faster to appropriate hyper-parameter values (Goodfellow *et al.* 2016). Finally, the user can run BioKEEN in *training mode* with the resulting hyper-parameter values.

To train the models, negative training examples are generated based on the algorithm described in Borders *et al..* To evaluate the trained models, BioKEEN computes two common evaluation metrics for KGE models: mean rank and hits@k.

## 3. Application

We used BioKEEN to train and evaluate several KGE models on the pathway mappings from ComPath (Domingo-Fernández *et al.*, 2018), the first manually curated intra- and inter-database pathway mapping resource that bridges the representations of similar biological pathways in different databases. Then, we used the best model to predict new relations representing pathway crosstalks and hierarchies. After removing reflexive triplets, we found that the highest ranked novel equivalence between TGF-beta Receptor Signaling (wikipathways:WP560) and TGF-beta signaling pathway (kegg:hsa04350) as well as the highest ranked hierarchical link that Lipoic acid (kegg:hsa00785) is a part of Lipid metabolism (reactome:R-HSA-556833) both represented novel crosstalks. Upon manual evaluation, each fulfilled the ComPath curation criteria and can be added to the resource.

We performed HPO for five different models to illustrate the need for choosing the appropriate hyper-parameter values. For the TransE model, comparing the hyper-parameters similar to those reported by Borders *et al.* with the hyper-parameters from HPO showed an improvement in the hits@10 metric from 19.10% to 63.20%.

Moreover, the nature of the model strongly influences the results. We found that the simpler models (e.g., TransE, UM, and DistMult) performed similar or even better than the more complex ones (e.g., TransH and TransR). This might be explained by the fact that the more expressive models overfit since ComPath is a not a large data set. Ultimately, this case scenario illustrates the ability of BioKEEN to assist users in finding reasonable combinations of models and their hyper-parameter values to predict novel links.

## 4. Discussion and Future Work

While BioKEEN already includes several models and components to build machine learning pipelines, it could still benefit from several additions and improvements. Because of the heterogeneity and lack of structure in most biological and clinical data, we aim to implement additional KGE models that incorporate text, logical rules, and images in addition to the triples in a knowledge graph (Wang *et al.,* 2017; Hamilton *et al.* 2018). Further, modeling multiscale biology (i.e., the *-omics*, biological process, phenotype, and population levels) results in KGs with a variety of compositions, structural features, and topologies for which different KGE models are most appropriate.

The current version of BioKEEN implements the negative sampling approach described by Bordes *et al.*, which could cause false negatives. One improvement could be to incorporating prior biological knowledge and constraints to generate triples guaranteed to be true negatives such as: i.) type constraints for predicates (e.g. the relation *transcribed* is only valid from gene to protein), ii.) valid attribute range for predicates (e.g., protein weight is below 1000 kDa) and iii.) functional constraints such as mutual exclusion (e.g., a protein is coded by one gene).

With BioKEEN, we have provided a software package that enables users without expert knowledge in machine learning to learn and apply biological based KGE.

## Supporting information

## Acknowledgements

We thank our partners from the Bio2Vec, MLwin, and SimpleML projects for their assistance.

## Funding

This research was supported by funding from King Abdullah University of Science and Technology (KAUST) in the Bio2Vec project (CRG6 grant 3454). *Conflict of Interest:* none declared.

