## Supplementary material for "BioKEEN: A library for learning and evaluating biological knowledge graph embeddings"

### Supplement

#### Outline

- **Supplementary Table S1:** Knowledge Graph Embedding Models implemented in BioKEEN
- **Supplementary Table S2**: Databases included in BioKEEN via Bio2BEL
- **Supplementary Table S3**: Hyper-Parameter optimization results on ComPath
- **Software Installation and Documentation**

| **Reference** | **Name** | **Description** |
|---|---|---|
| Bordes, *et al.* [1] | TransE | Considers a relation as a translation from the head to the tail entity. |
| Wang, *et al*. [2] | TransH | Extends TransE by applying the translation from head to tail entity in a relational-specific hyperplane. |
| Lin, *et al* [3] | TransR | Extends TransE and TransH by considering different vector spaces for entities and relations. |
| Ji, *et al.* [4] | TransD | Extends TransR to use fewer parameters. |
| Dettmers, *et al.* [5] | ConvE | Uses a convolutional neural network (CNN) for applying link prediction. An input instance is represented by the subject and predicate embeddings, these are rescaled to represent an “image”, and on the rescaled input the CNN is applied. The CNN will predict the most probable object. |
| Bordes, *et al.* [6] | SE | For each relation head and tail entity are projected by different matrices. |
| Bordes, *et al.* [7] | UM | Simplifies TransE by ignoring relation embeddings. |
| Nickel, *et al.* [8] | RESCAL | Represents relations as matrices and models interactions between latent features. |
| Dong, *et al.* [9] | ERMLP | Neural network based approach in which subject, predicate and object are concatenated and fed to the neural network. |
| Yang, *et al.* [10] | DistMult | Simplifies RESCAL by restricting matrices representing relations as diagonal matrices. |

**Supplementary Table S1**. Knowledge graph embedding models implemented in BioKEEN.

| Reference | Database | Type | Bio2BEL Zenodo |
|---|---|---|---|
| [11, 12] | ADEPTUS | disease-differential expressed genes | [13] |
| [14] | ComPath | pathway-pathway | [15] |
| [16] | DrugBank | drug-target | [17] |
| [18] | ExPASy | protein-enzyme class | [19] |
| [20] | HIPPIE | protein-protein interaction | [21] |
| [12, 22] | HSDN | disease-symptoms | [23] |
| [24] | KEGG | protein-pathway | [25] |
| [26] | mirTarBase | miRNA-target | [27] |
| [28] | MSigDB | protein-pathway | [29] |
| [30] | Reactome | protein-pathway | [31] |
| [32] | InterPro | protein-domains and protein-family | [33] |
| [34] | WikiPathways | protein-pathway | [35] |

**Supplementary Table S2**. Biological databases included in BioKEEN via Bio2BEL until the date.

| Model | EED | RED | LR | Loss Function | Margin | Normalization | Scoring Function | Batch Size | Epochs | WSC | FCT | Seed | Mean Rank | Hits@10 (%) |
|---|---|---|---|---|---|---|---|---|---|---|---|---|---|---|
| TransE | 150 | - | 0.01 | MRL | 12.0 | L2 | L1 | 32 | 2500 | - | Yes | 2 | **131.44** | **63.20** |
| | 50 | - | 0.01 | MRL | 5.0 | L2 | L1 | 32 | 2200 | - | Yes | 2 | 130.99 | 62.08 |
| | 50 | - | 0.01 | MRL | 1.0 | L2 | L1 | 32 | 1000 | - | Yes | 2 | 226.03 | 19.10 |
| TransH | 50 | - | 0.01 | MRL | 4.0 | - | L2 | 400 | 1500 | 0.03 | Yes | 2 | 481.08 | 25.00 |
| TransR | 30 | 50 | 0.01 | MRL | 0.5 | - | L1 | 32 | 1500 | - | Yes | 2 | 200.13 | 41.01 |
| DistMult | 150 | - | 0.01 | MRL | 15.0 | - | - | 32 | 2000 | - | Yes | 2 | 230.33 | 38.48 |
| UM | 200 | - | 0.01 | MRL | 15.0 | L2 | L2 | 64 | 1000 | - | Yes | 2 | 224.83 | 43.26 |

**Supplementary Table S3**. Hyper-Parameter optimization results of TransE, TransH, TransR, DistMult, and UM evaluations on ComPath. We used a random 90:10 split for splitting the dataset into a training and test set. Acronyms: EED (Number of Entity Embedding Dimensions), FCT (Filtering Corrupted Triples), LR (Learning Rate), and MRL (Margin Ranking Loss), RED (Number of Relation Embedding Dimensions), WSC (Weight value for Soft Constraints). Different parameter settings are presented for TransE to illustrate the sensitivity of choosing appropriate hyper-parameter values.

#### Software Installation and Documentation

All packages described in the manuscript are available through GitHub (https://github.com/) or PyPI (https://pypi.org), the main packaging system for Python 3, under the MIT license. All relevant information for installation is bundled in the package, so it can be easily and quickly installed independently of the operating system, running any modern version of the Python programming language. The documentation for all packages was built using the Python documenting tool Sphinx and is accessible at Read the Docs (https://pykeen.readthedocs.io/ and https://biokeen.readthedocs.io/). Releases to PyKEEN and BioKEEN are also tracked by Zenodo (https://zenodo.org) under [36] and [37], respectively.
